# Simulating digital micromirror devices for patterning coherent excitation light in structured illumination microscopy

**DOI:** 10.1101/2020.10.02.323527

**Authors:** Mario Lachetta, Hauke Sandmeyer, Alice Sandmeyer, Jan Schulte am Esch, Thomas Huser, Marcel Müller

## Abstract

Digital micromirror devices (DMDs) are spatial light modulators that employ the electro-mechanical movement of miniaturized mirrors to steer and thus modulate the light reflected of a mirror array. Their wide availability, low cost and high speed make them a popular choice both in consumer electronics such as video projectors, and scientific applications such as microscopy.

High-end fluorescence microscopy systems typically employ laser light sources, which by their nature provide coherent excitation light. In super-resolution microscopy applications that use light modulation, most notably structured illumination microscopy (SIM), the coherent nature of the excitation light becomes a requirement to achieve optimal interference pattern contrast. The universal combination of DMDs and coherent light sources, especially when working with multiple different wavelengths, is unfortunately not straight forward. The substructure of the tilted micromirror array gives rise to a *blazed grating*, which has to be understood and which must be taken into account when designing a DMD-based illumination system.

Here, we present a set of simulation frameworks that explore the use of DMDs in conjunction with coherent light sources, motivated by their application in SIM, but which are generalizable to other light patterning applications. This framework provides all the tools to explore and compute DMD-based diffraction effects and to simulate possible system alignment configurations computationally, which simplifies the system design process and provides guidance for setting up DMD-based microscopes.

## 1. Introduction

Spatial light modulators (SLMs) offer a robust and fast way to pattern the excitation light in a fluorescence microscope. This can be employed for various illumination schemes [1], for example to achieve selective (de)activation of photo-switchable dyes [2], and most notably, to achieve background suppression and resolution enhancement in *structured illumination microscopy* (*SIM*) [3–8], a widely used, fast super-resolution microscopy technique [9–13]. Typical SLMs are based on liquid crystal technology, and thus operate by electrically modulating the phase (or amplitude) of light through their active material. Digital micromirror devices (DMDs), on the other hand, work electro-mechanically, by individually flipping the orientation of each mirror between 2 pre-defined states. Because of their widespread use in consumer devices such as video projectors, DMDs are available at relatively low cost and in a variety of sizes. They also offer high switching speeds, they can handle high light intensities, and depending on coating, are not sensitive to light polarization. This makes them an interesting option for many SLM applications in microscopy [14–22]. However, the jagged nature of the micromirror array gives rise to the *blazed grating effect* that becomes rather annoying and detrimental when using DMDs in combination with a coherent light source [23,24]. Thus, if DMDs are selected as active light modulation systems in a fluorescence microscope based on laser light sources, this effect must be well understood and needs to be taken into consideration.

The work presented here was motivated by our wish to expand the range of applications of a DMD-based SIM system (see Figure 1), where a coherent laser source is the primary source for the SIM interference pattern, giving rise to optimal pattern contrast [14]. We have developed a set of simulation frameworks, that allow us to simulate the propagation of coherent light reflected off of a DMD at different angles of incidence, at different wavelengths and with the DMD displaying arbitrary patterns. This facilitates us to explore the feasibility of optical layouts, to determine which simplifications (e.g. keeping optical elements in a single plane on a table) are possible, and to choose the proper wavelength combinations that could be used in a multi-color DMD-based system. While our work is motivated by and centred around SIM, most of the findings should apply to any other, arbitrary DMD-based microscope systems that employ coherent illumination.

**Figure 1:**
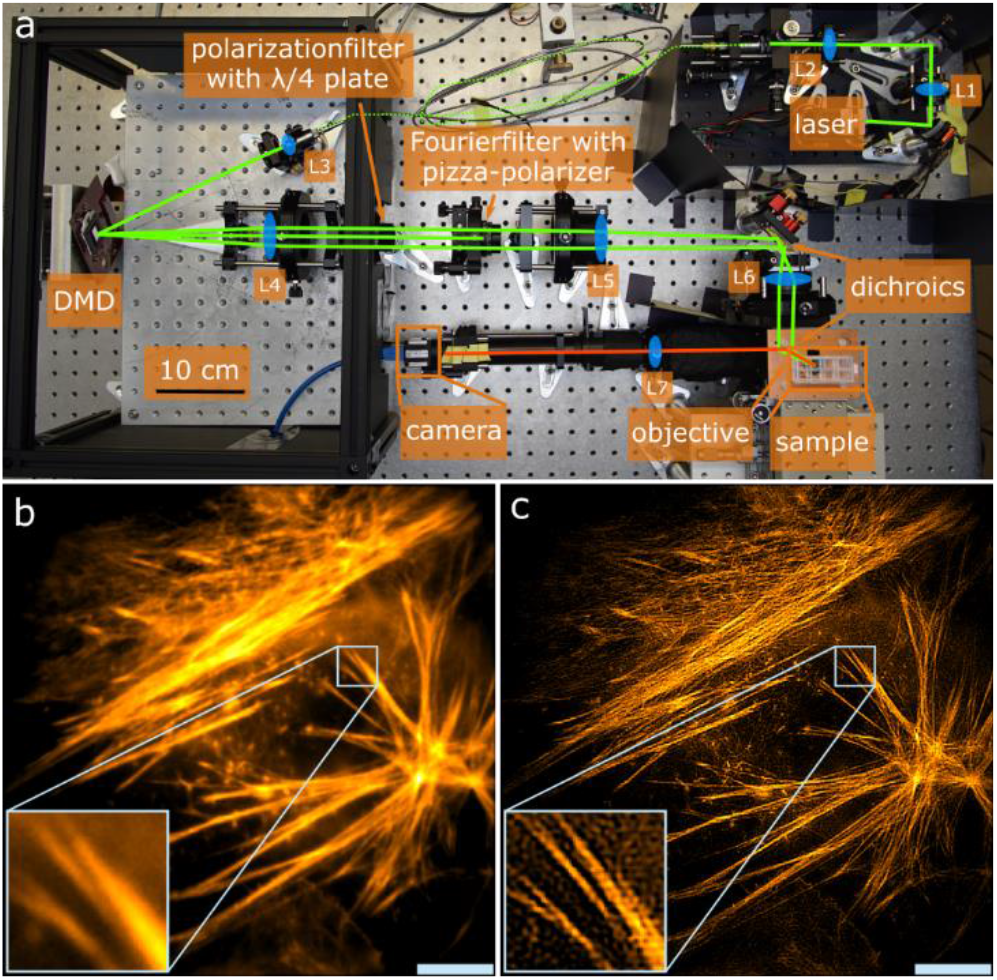
DMD-SIMmicroscope and measurements. (a) Compact & cost-effective SIM system based on a DMD with a 532 nm laser. The design of this microscope necessitated and motivated the research presented in this article. (b) 36 μm x 36 μm wide field of fixed U2OS cell labeled with Phalloidin Atto532 acquired using the instrument with 20 ms exposure per raw frame. The actin filaments are not distinguishable. (c) SIM data of (b). The actin filaments are distinguishable. (scale bar 5 μm, inset 2.8 μm x 2.8 μm)

## 2. Methods

If DMDs are to be used with coherent illumination, it is essential to know and understand the resulting diffraction patterns generated by both the pattern displayed on the DMD and, much more importantly, the structure of the tilted micromirrors that is native to the device. For this purpose, we have developed three simulation algorithms, each with different assumptions and resulting strengths. Additionally, we have developed an analytical solution that can be used along the diagonal of the DMD. All three algorithms are based on the same physical model of propagating electromagnetic fields interacting with the DMD’s structure.

### General modelling of coherent light and DMDs

The general model was derived for the a previously developed DMD-based SIM system [14], and, for completeness, is summarized here, as it forms the basis of our simulation approaches. For incident and diffracted directions we use the normalized vectors 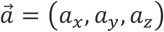 and 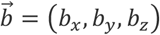 with the angle coordinates (*φ_a_, ϑ_a_*) and (*φ_b_, ϑ_b_*):

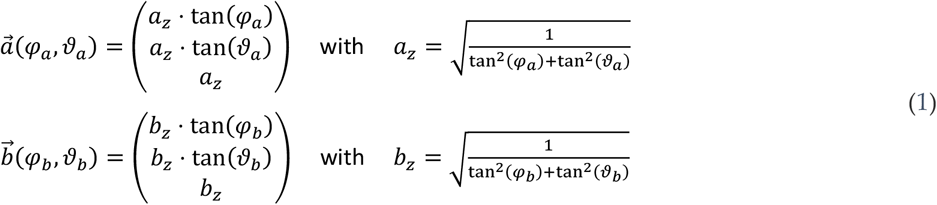

For every incident direction, all diffracted directions must be considered. Since the DMD is basically a two-dimensional array of mirrors, we start with modelling a single mirror with defined dimensions:

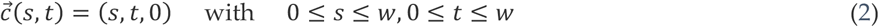

Here, *w* is the width and height of the mirror, and *s* and *t* are parameters for the x- and y-directions, respectively. To model the tilt state each single mirror in a DMD array can be rotated by the angle *γ* around the normalized diagonal axis 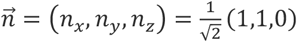 via the rotation matrix:

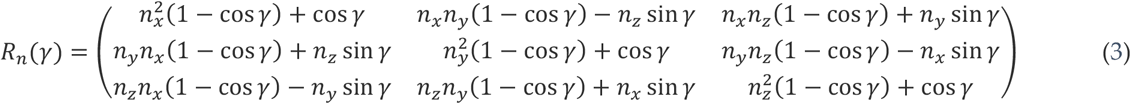

To shift a single mirror, we use the native grid 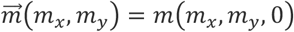 with the micromirror pitch *m* = *w* + *g*. The gap between the non-tilted mirrors is *g*. 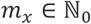 and 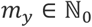 are the indices of the grid points (the single mirrors). This leads us to the following expression to describe each single mirror on the DMD:

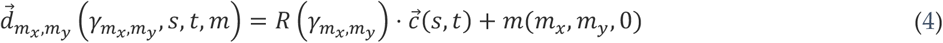

We can model monochromatic coherent light using the Fraunhofer / far-field approximation with a time-independent electric field, which depends on the beam profile 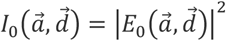 projected onto the DMD:

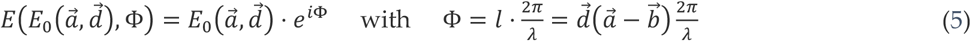

*E*_o_ represents the incidence amplitude and Φ the resulting phase, which is dependent on the path length 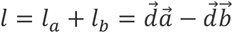 and the wavelength *λ* (Figure 2a). For the electric field diffracted at a specific point on the DMD we obtain:

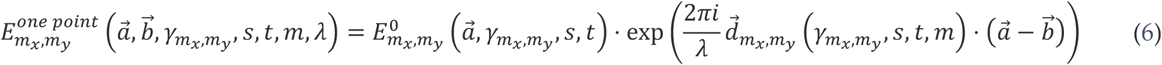

**Figure 2:**
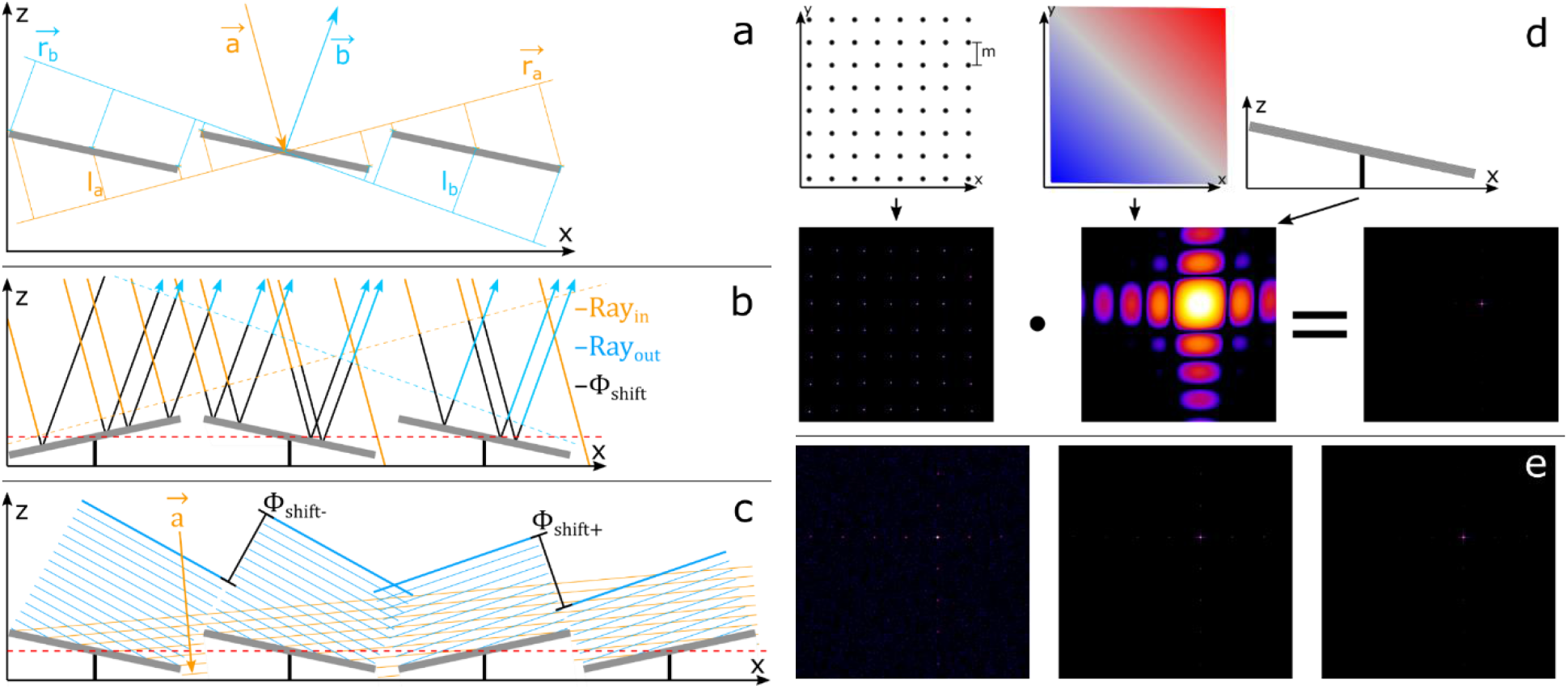
Illustration and comparison of algorithms for the simulation of diffraction with DMDs. Modelling of (coherent) light that is diffracted at a DMD surface. All diffraction images shown are in logarithmic intensity representation and were generated with an array of50 x 50 micromirrors at 532 nm wavelength. Each micromirror has a size of 7.56 μm x 7.56 μm. φ_a_ = –ϑ_a_ = –21° (blazed condition) was chosen as angle of incidence. The diffraction images are shown for both angles φ_b_ and ϑ_b_ with an angle range of - 15° to 15°. (a) General determination of the phase shift for different points of a planar wave front, which is incident on the DMD in direction 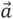 and is diffracted in direction 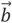. This approach is correct in the far-field / Fraunhofer approximation. (b) Ray tracing approach: Modeling of rays and their phase shift, which are incident on the DMD in the direction 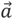 and diffracted in the direction 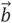. (c) Analytical phase shift approach: Simplified modelling of diffraction images in the form of wave fronts reflected by micromirrors, where the diffraction image of a single mirror is analytically known. (d) Grating approach: In the upper row the native grating of the DMD array, and a single mirror are shown schematically. In the lower row the corresponding diffraction images and their product, which results in the diffraction image of the entire DMD, are shown. (e) Comparison of the diffraction images (from left to right): Ray tracing approach, analytical phase shift approach, grating approach.

To calculate the diffracted field distribution we need to integrate over each single mirror and calculate the sum over the entire DMD with *N_x_* and *N_y_* as the number of single mirrors in x- and y-direction:

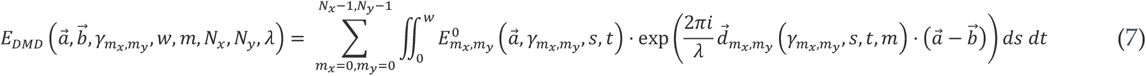

This expression depends on many parameters and generally cannot be simplified further analytically. In order to be able to calculate diffraction patterns numerically and in adequate time, it is thus necessary to make further assumptions to simplify the calculation. Depending on how these assumptions are chosen, different approaches emerge capable of modelling different system constraints:

### Ray tracing approach

We approximate integrating over every single mirror and summing over the DMD (eq. (*7*)) by running a Monte-Carlo-simulation with rays which will be summed up (Figure 2b). This yields a computationally feasible (if somewhat slow) simulation that does not introduce further constraints or approximations not inherent to the Monte-Carlo process. To simulate a collimated Gaussian incident laser beam, we use a Gaussian probability density for the generation of *K* randomized incident beams with this equation of a line

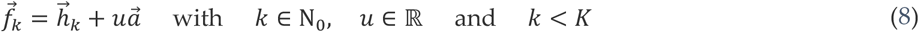

and with 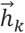 as support vector, 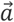 as normalized incident direction vector and the parameter *u*. Here, 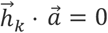 because 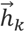 and 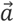 are perpendicular to each other. 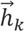 is Gaussian randomized for each ray 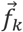. To obtain the point of diffraction on the DMD we must calculate the intersection point 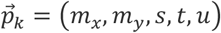 of 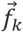 and 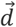. For each ray we assume that the intensity is 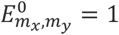. This simplifies eq. (*6*) to:

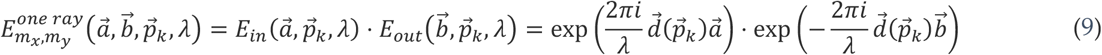

A Gaussian beam profile is already considered by the probability distribution of the rays. The field composed of contributions from all rays needs to be summed up to result in the final ray tracing expression for the diffracted field distribution (Figure 2e, left). This simplifies eq. (*7*) to:

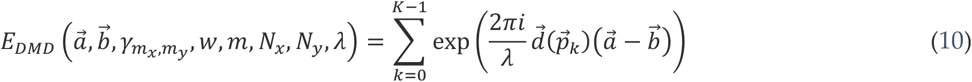

In this ray-tracing approach, both the assumption of a Gaussian beam profile and especially of fully monochromatic (i.e. coherent) light could easily be changed. Thus, by using a different ray distribution, and by allowing for the single rays (eq. 9) to follow a wavelength and phase distribution when being summed up (eq. 10), it would easily be possible to simulate for example an arbitrary profile of a partially coherent source.

### Analytic phase shifting approach

For this approach we assume that the field amplitude over each single mirror is constant: 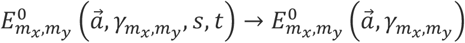 and each single mirror can only be in the tilt state *γ*^−^ or *γ*^+^ (as reasonable assumption for the steady state of a DMD). A Gaussian beam profile can still be approximated via varying 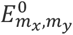 over the different single mirrors, and as typically a reasonably sized array of mirrors is illuminated, the error introduced by this approximation is small. This provides us with the opportunity to solve the integral in eq. (*7*) over a single mirror for *γ*^−^ and *γ*^+^, with the analytically known solution for 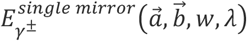 (see dmd.nb in the attached repository). Instead of calculating the field for each mirror individually, we can use the fields of a reference mirror for *γ*^−^ and *γ*^+^ and apply the phase shift for the desired grid point (*m_x_, m_y_*) shown in Figure 2c. This simplifies eq. (*7*) for the diffracted field (Figure 2e, middle) to:

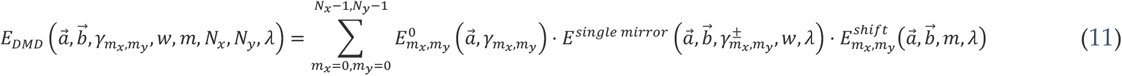

This approach is computationally much simpler, while only introducing a minor approximation. It was thus used for the design of our compact DMD-based SIM system [14].

### Grating approach

Additionally to the assumptions for the analytic phase shifting approach, we assume here that the field amplitude over the entire DMD is constant 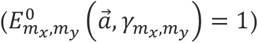 and that all single mirrors are in the *γ*^−^ or *γ*^+^ state. The field 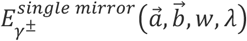 can be considered as envelope over the diffracted native DMD grid 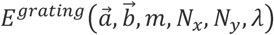. This gives the opportunity to write eq. (*7*) as follows:

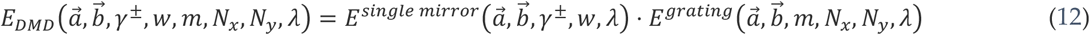

To get the intensity *I_DMD_* we calculate and multiply both diffraction patterns *I_envelope_* and *I_grating_* (Figure 2d), very similar to the case of the Young double slit experiment:

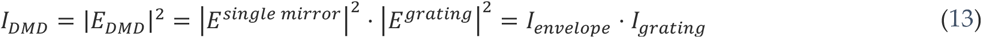

Of course, this approach is constrained to the simulation of a DMD without a pattern being displayed. However, if only the diffractive nature of the DMD itself is of concern, this offers a computationally very effective solution.

### Blaze condition approach

We assume that the incident beam is perpendicular to the tilting axes 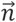 of the single mirrors. This leads to 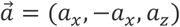 and –*φ_a_* = *ϑ_a_*. To imagine it simply we rotate the DMD by 45° (Figure 3a). To account for the rotation we switch to new angle coordinates (Figure 3b):

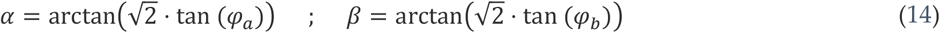

**Figure 3:**
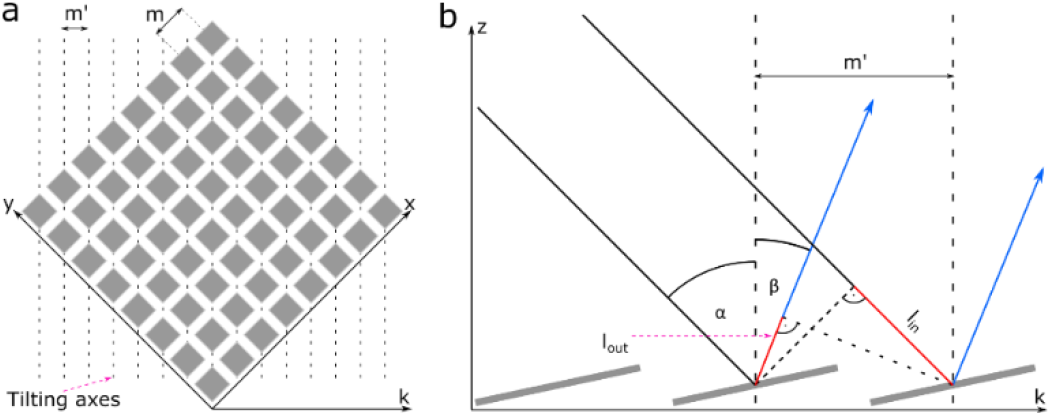
Blazed condition approach. (a) Schematic illustration of a DMD rotated by 45° around the z-axis. Furthermore, the tilt axes of the micromirrors, the grid constants *m* and *m′* and the coordinate axis *k* are shown. (b) Graphical representation of the calculation of the phase shift for light, which is incident along the KZ-plane on a DMD.

As new coordinate axes we define 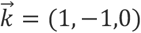. Along 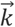 we get a new lattice constant 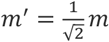.

The pathlength is given by

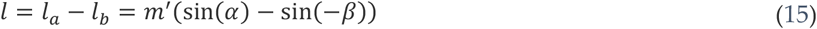

For perfect constructive grating interference, the path difference *l* must be an integer multiple of the wavelength *λ*:

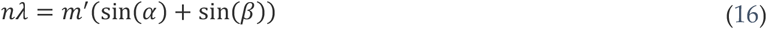

If we have perfect constructive interference in the center of the envelope, the blaze condition is fulfilled. The center of the envelope can then be assumed as a reflection of the light incident on the surface of a single mirror and we define this as our diffraction angle *β* = –*α* + 2*γ*. For this reflected envelope we can now calculate the corresponding diffraction order:

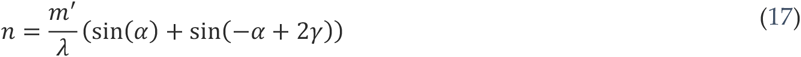

If *n* is an integer the blaze condition is fulfilled. We use *v* to visualize this relationship:

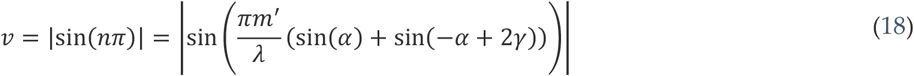

Here, *v* = 0, whenever the blaze condition is fulfilled. This allows us to visualize the blaze condition for different wavelengths *λ* and different incident angles *α* along the diagonal of the DMD (Figure 3b). While this approach is even more constrained, it has an important real-world application: It allows us to directly calculate incident and reflective angles when placing a DMD rotated by 45° on an optical table, where collimated beams are expected to run parallel to the table’s surface. Here, it provides a quick solution for the instrument design. A similar approach was pursued by Li et al. [25]

## 3. Results & Discussion

Each of the algorithms described above has certain advantages and disadvantages, which means that each algorithm has its own area of application. As shown in figure 1e, the ray-tracing approach, the analytic phase shifting approach and the lattice approach each deliver qualitatively the same results under identical boundary conditions. The positions of the individual diffraction orders are identical in the three algorithms. All diffraction images shown in this article were simulated with *m* = 7.56 μ*m* micromirror pitch and *γ*^±^ = ±12° as tilt angle, which corresponds to the dimensions of the *DLP*^®^ *LightCrafter*^™^ *6500* (*Texas Instruments*) [26] and some other DMDs. To obtain a clear visualization of the diffraction orders with large visible spots, the simulation is limited to an array of 50×50 mirrors being illuminated. Of course, when no or a different visualization is needed, the algorithms can be run for larger mirror arrays.

The ray tracing approach is most flexible because it requires the least simplifying approximations. It can be used to calculate the diffraction patterns for any DMD-pattern. In addition, the approach could be extended relatively easily for arbitrary exposure beam profiles and incoherent light by specifying a corresponding spatial distribution and wavelength distribution and/or phase distribution for the random rays. The main disadvantage of the ray tracing approach is that it is by far the slowest of the presented algorithms. Therefore, this algorithm is especially suitable for the simulation of incoherent or partially coherent light. The analytical phase shifter has the only disadvantage that, when compared to the ray tracing approach, it cannot be readily modified for use with incoherent illumination, or if done so, it would lose its speed advantage. But the algorithm works much faster and is especially suitable for the calculation of diffraction images where the DMD is provided with a pattern (see Figure 4).

**Figure 4:**
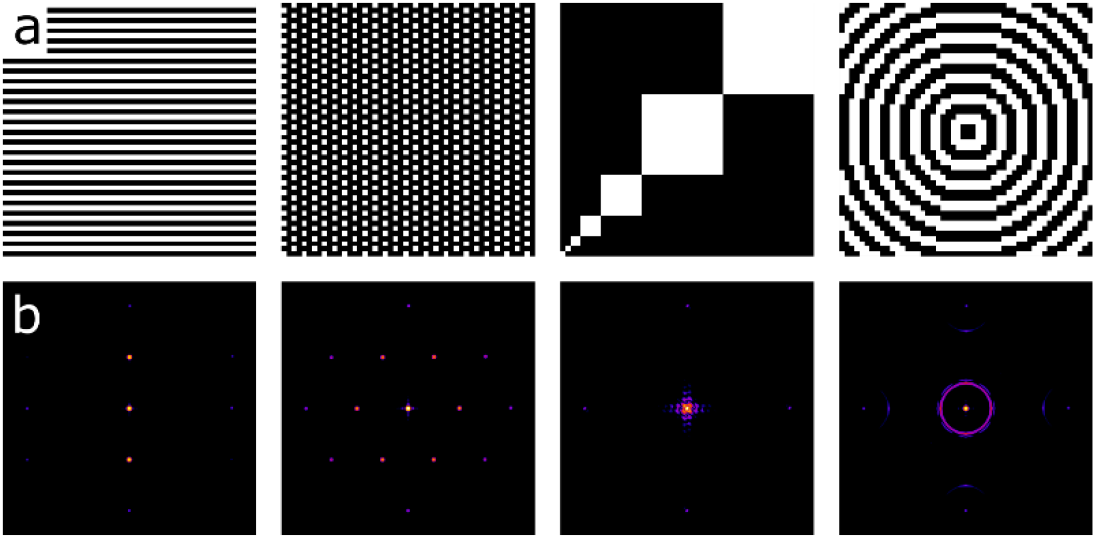
Comparison of diffraction images by different patterns generated on the DMD. (a) Different DMD patterns. From left to right: Horizontal lines; MAP-SIM example; MAP-SIM calibration pattern [30–32]; concentric circles. (b) Diffraction images of the DMD patterns shown in (a), simulated with the analytical phase shift approach with 50 x 50 micromirrors at 532 nm wavelength. For the angles of incidence *φ_a_* = –ϑ_a_ = –21° (blaze condition) was chosen. The diffraction images are shown for a range of - 1.8° to 8.2° for *φ_b_* (x axis) and −8.2° to 1.8° for *ϑ_b_* (y axis). The intensity distribution is shown on a logarithmic scale.

The grating approach is not able to simulate any patterns displayed on the DMD. It assumes that all mirrors are tilted in the same direction. Due to its high speed, however, the algorithm is very well suited for investigating the effects of different boundary conditions such as changes in wavelength and angle of incidence on the diffraction image generated natively by the structure of the micromirror array (Figure 5). It can be used to determine the distance between the center of the envelope and the brightest native grating order (Figure 5a). If this distance is 0°, the blaze condition (Littrow configuration) is fulfilled. For the use of coherent light in e.g. a SIM microscope, this is exactly what is desired, because then an isotropic envelope field and intensity distribution is present in the Fourier plane. This has the consequence that the disadvantages of the blazed grating effect of DMDs can be negated by exploiting the blaze condition. This is true for a range of angles of incidence which are shown in black in Figure 5b. These angles of incidence that fulfil the blaze condition then result in the angles of diffraction marked in Figure 5c.

**Figure 5:**
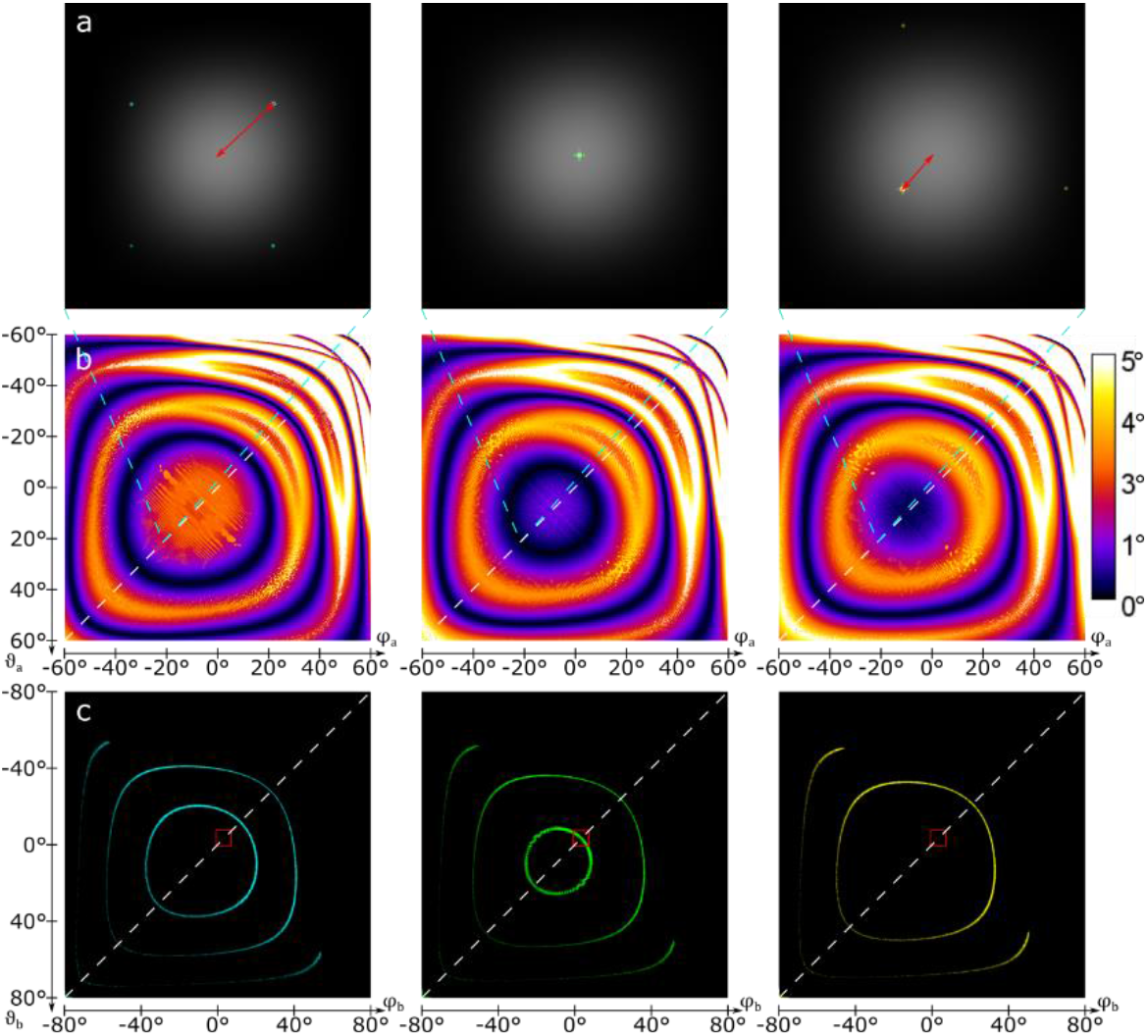
Analysis of the displacement between the envelope and the brightest diffraction order of the DMD. The columns from left to right correspond to the wavelengths 488 nm, 532 nm and 561 nm. Row (a) illustrates the displacement between the center of the envelope (gray, linear intensity representation) and the brightest diffraction order of the native DMD grating (cyan, green, yellow, logarithmic intensity representation) using red arrows. The field of view ranges from −1° to 7° for *φ_b_* and −7° bis 1° for *ϑ_b_* at an angle of incidence of *φ_a_* = –ϑ_a_ = –21°. Row (b) visualizes the offset which can be determined from (a) for the angles of incidence *φ_a_* and *ϑ_a_* ranging from −60° to 60°. The dark areas with a displacement close to 0° indicate angles which fulfill a blaze condition and are well suited for illumination with coherent light. The cyan dotted lines between (a) and (b) indicate an angle of incidence of *φ_a_* = –ϑ_a_ = –21° for the diffraction images seen in (a). Row (c) shows the areas in the diffraction space of *φ_b_* and *ϑ_b_* from −80° to 80°, each, in which the displacement between the center of the envelope and the brightest diffraction order of the DMD is not more than 0.1°. Therefore, angles of incidence in a range of −60° to 60° used in (b) were used for *φ_a_* and *ϑ_a_*. The red boxes mark the diffraction space shown in (a). The white dotted lines in (b) and (c) correspond to the diagonal angles of incidence that are considered in the blazed condition approach.

If a DMD-based SIM microscope should require the use of multiple excitation wavelengths, then it is necessary that for each wavelength the blaze condition is fulfilled. A possible approach to solving this problem is to use the *γ*^−^ position of the micromirrors for one wavelength and the *γ*^+^ position for the other wavelength. As shown for example for 488 nm or 638 nm in combintion with 561 nm in Figure 7a, there are two configurations in which the blaze condition is fulfilled for both wavelengths. This configuration is, however, very difficult to achieve experimentally (as multiple independent tip/tilt axis have to be precisely aligned) and it is also very susceptible to the slightest change in angle of the micromirrors (we found e.g. 0.3° in [14]). Therefore, we propose to limit the angle of incidence and thus also the outgoing angle for the optical axis of the DMD-SIM microscope to the diagonal perpendicular to the tilting axis of the micromirrors. This corresponds to a rotation of the DMD of 45° around the z-axis (Figure 3). This limits the possible angles of incidence as well as the angles of diffraction for the optical axis to the white dotted diagonal in fig 5b. This reduces the number of degrees of freedom of the incident laser beam (*φ_a_, ϑ_a_*) → *α* and the optical axis (*φ_b_, ϑ_b_*) → *β* from two to one each, as described in eq. (*14*). The experimental implementation thus becomes much easier, because the exposure path of the microscope can be set up parallel to the optical table, as usual. Now, however, the wavelengths must be chosen skilfully. For a more precise analysis of the blaze condition along the diagonal, both the grating approach and the blazed condition approach are suitable.

**Figure 6:**
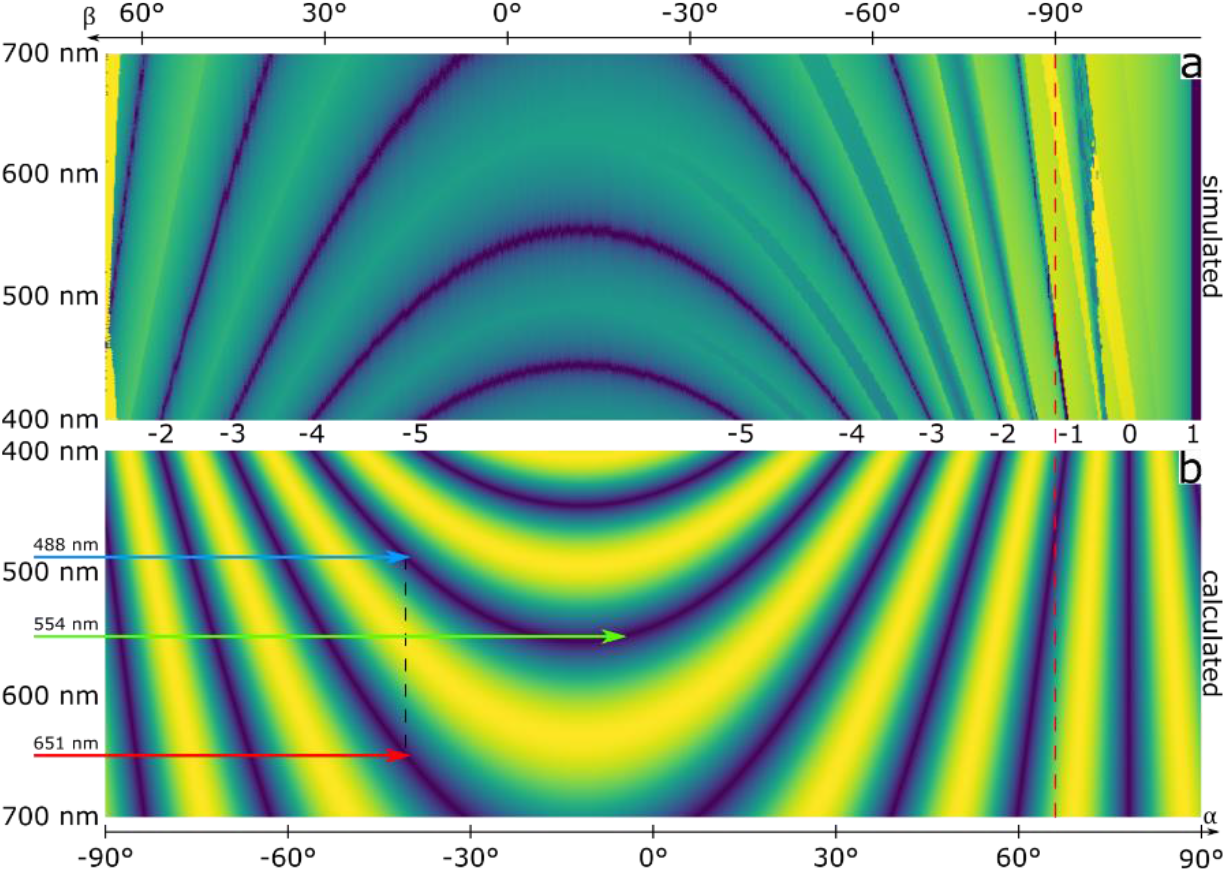
Blaze condition along the diagonals shown in Figure 3b. Visualization of the blaze condition along the diagonals shown in Figure 3b for the visible spectrum. In the dark areas the blaze condition can be considered fulfilled. For orientation, the wavelength range is shown on the left-hand side. The angles of incidence and diffraction are shown at the top and the bottom of the graph. The vertical red dashed line marks the diffraction angle of 90°. Angles above 90° are irrelevant for practical applications. (a) The blaze condition is measured by the distance between the center of the envelope and the brightest diffraction order of the DMD, which was simulated by the grating approach. The measured distances are shown as logarithmic intensity distribution. (b) Visualization *v* = |sin(*nπ*)| of the blaze condition using the blazed-grating approach. The integer diffraction orders belonging to the blaze conditions are shown in the middle between (a) and (b). The blue, red and green arrows point to an exemplary three-color combination for a DMD-SIM microscope with 488 *nm*, 651 *nm* and 554 *nm*. The black vertical dotted line is intended to illustrate that 488 *nm* and 651 *nm* fall on the DMD at the same angle of incidence.

**Figure 7:**
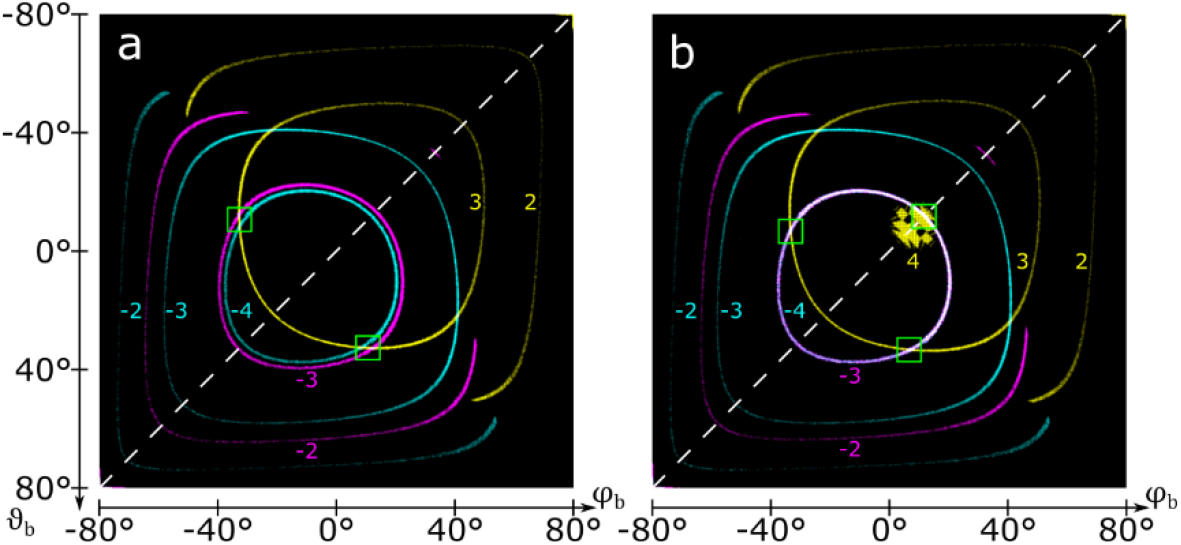
Multi color diffraction angles with blaze condition. Areas in the diffraction space for *φ_b_* and *ϑ_b_* of −80° to 80°, each, in which the displacement between the center of the envelope and the brightest diffraction order of the DMD is not more than 0.1°. Therefore, angles of incidence in a range of −60° to 60° were used for *φ_a_* and *ϑ_a_*. The white dotted lines correspond to the diagonal angles of incidence that are considered in the blazed condition approach. The colored numbers near the ring structures represent the corresponding diffraction orders. (a) Typical wavelengths 488 *nm* (displayed in cyan) & 638 *nm* (displayed in magenta) with *γ*^−^ = –12° and 561 *nm* (displayed in yellow) with *γ*^+^ = 12° are not well suited for multicolor applications. The green boxes mark possible configurations for two color configurations of 488 *nm* & 561 *nm* or 638 *nm* & 561 *nm*. (b) Optimized multicolor wavelengths 488 *nm* (cyan) & 651 *nm* (magenta) with *γ*^−^ = –12° and 554 *nm* (yellow) with *γ*^+^ = 12°. The green boxes mark possible configurations for three color configurations.

Both approaches show qualitatively the same results, therefore it is sufficient to refer to the blaze condition approach in the following. It is possible to use two wavelengths *λ*_1_ and *λ*_2_ with the same angle of incidence *α*_1,2_ and diffraction *β* with the same tilt state *γ*^−^ = –12°. Starting from eq. (*16*) it follows that

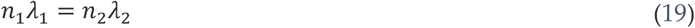

As integer diffraction orders *n*_1_ = –4 and *n*_2_ = –3 are suitable for the visible range. If we pretend *λ*_1_ = 488 *nm* we obtain:

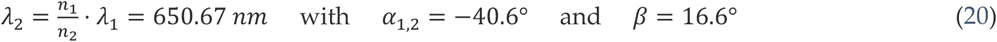

If a third excitation color is supposed to utilize the other tilt state *γ*^+^ = 12°, we get *α*_3_ = –*β* + 2*γ*^+^ = 7,4° as incidence angle and must assume *n*_3_ = 4 to be in the visible range, which then limits the wavelength to:

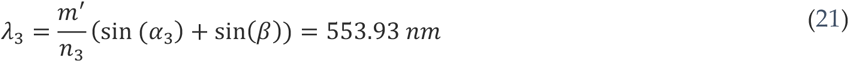

This combination of these three wavelengths, arranged at (–40.6°, 488 *nm*), (–40.6°,651 *nm*) and (7.4°, 554 *nm*) is indicated in Figure 6 as a possible configuration for a three color DMD-SIM microscope. Given that these wavelengths are close to the typical combinations of blue (488nm), yellow (561nm, 568nm) and red (635nm to 650nm) laser excitation sources used in fluorescence microscopy, and that suitable diode laser sources (even offering temperature-based finetuning of their emission wavelength) are readily available, this combination should make for a surprisingly capable three-color DMD-based microscope.

Further possible configurations are e.g. 473 *nm*, 631 *nm* and 551 *nm* or 491 *nm*, 655 *nm* and 555 *nm*. It is important to note here that *λ*_3_ depends very much on the tilt angles of the micromirrors, and thus might have to be adjusted based on the parameters of the specific DMD device. For and *λ*_2_ the tilt angle is of less importance. Looking at the diffraction space in which the blaze condition is fulfilled for typical wavelengths (Figure 7a), it is noticeable that, using the same tilt state, no matches are found. The ringlike structures in Figure 7 represent the different diffraction orders. Their size depends on wavelength and diffraction order and for clarity they are shown in different colors. As mentioned earlier, using the other tilt state results in matches, which are not on the diagonal and therefore not suitable for experiments. But if we choose the wavelengths carefully for multicolor applications (fig 7b), the circular structure of the −4th and −3rd diffraction order of 488 nm and 651 nm using the same tilt state match exactly. If we use the other tilt state for 554 nm, there is, besides the matches outside the diagonal, another one on the diagonal, which is formed by a collapsing/emerging circular structure of the 4th diffraction order. This configuration, already described above, is therefore ideal for a three-color DMD-SIM microscope.

## 4. Conclusions

The framework and the software tools provided here provide a rather universal set of simulations tools for experimentalists planning to use DMDs as spatial light modulators with coherent light sources in SIM microscopes or similar applications. Compared to ferro-electric light modulators (FLCoS-SLM), DMDs are faster and more cost efficient, and they allow for a more compact design due to the smaller micromirror/pixel pitch. Also, the use of FLCoS-SLMs requires the implementation of a much more complicated timing scheme because of the limited duration during which a specific pattern can be projected by these devices. DMDs have no such constraints and can switch at speeds up to an order of magnitude faster than FLCoS-SLMs. We have developed four different approaches to simulate coherent light diffracted and reflected by a DMD (table 1). The results provide practical design suggestions for the circumventing the undesired blazed-grating effect of DMDs. Therefore, it is now possible to design a multicolor DMD-based SIM microscope with simple opto-mechanics which can fully utilize all of the advantages of a DMD mentioned before. We are currently building a two-color DMD SIM microscope and plan to add a third color to it in its final implementation.

**Table 1:** Comparison of all discussed approaches.

| Approach | Ray tracing | Analytic phase shifting | Grating | Blaze condition |
| --- | --- | --- | --- | --- |
| Capable of patterns | + Yes | + Yes | - No | - No |
| Speed (Time CPU/GPU) | -- Very slow (84 min/-----) | - Slow (7.5 min/3 sec) | + Fast (3.2 sec/1.2 sec) | ++ Instant |
| Use case | Incoherent light | Specific DMD-patterns | Native pattern under different conditions | Analysis of the diagonal |

This table gives a compact summary of the discussed approaches. The runtimes for the approaches were measured with 50 x 50 mirrors at a resolution of 1500 x 1500 pixels in diffraction space on an Intel Core i5 4690 4x 3.50GHz with a NVIDIA GeForce GTX 1060 6GB. The ray tracing approach was measured with 100000 rays. The runtimes can vary significantly depending on the approach, the number of mirrors, the resolution and the computer used.

The results shown in this article can easily be reproduced with the ImageJ/Fiji plugin which we provide as an open access download. The plugin contains all approaches and algorithms presented in this article, which can be adapted to different system conditions. For a more detailed insight into the plugin and some mathematical additions we recommend a look at the supplementary material.

Following the spirit of open science, we provide all source code and raw data for the results presented in this manuscript to the scientific community. All code is openly accessible under GPLv2 (or later) license and can be found in online repositories under http://github.com/fairSIM and github.com/biophotonics-bielefeld.

## Additional Information

### Data Accessibility

A software package to simulate with the introduced algorithms the diffraction patterns of DMDs and the introduced further analysis depending on the angle of incidence, light wavelength and DMD pattern has been created. The software is written in Java to work as a plugin to scientific image processing package ImageJ/FIJI [27,28]. and provides GPU acceleration through the jCuda framework [29]. The source code is available under an open-source (GPLv3) licence in an online repository: github.com/biophotonics-bielefeld/coherent-dmd-sim-simulator.

### Authors’ Contributions

ML wrote the DMD simulation software, developed the analytic phase shifting approach, the grating approach, the blazed condition approach, set up electronic and control software of the DMD-SIM microscope and drafted the manuscript. HS developed the ray tracing approach and crosschecked math and simulations. AS built the DMD-SIM microscope, performed the SIM measurements and processed the SIM reconstructions. TH and JSaE supervised the project and helped write the manuscript. MM envisioned the project and wrote the manuscript. All authors read and approved the manuscript.

### Competing Interests

The authors declare that they have no competing interests.

### Funding Statement

We acknowledge funding by the Deutsche Forschungsgemeinschaft (DFG, German Science Foundation) - project number 415832635. This project has also received funding from the European Union’s Horizon 2020 research and innovation programme under the Marie Skłodowska-Curie grant agreements No. 752080. ML was supported by the Protestant Hospital of Bethel Foundation.

## Notes

### Competing Interest Statement

The authors have declared no competing interest.

