## Supplementary information for "Simulating digital micromirror devices for patterning coherent excitation light in structured illumination microscopy"

### 1. Mathematical Supplementals

**Raytracing Approach:**

The intersection calculation of the individual rays with the DMD can be described as follows. To get the point of diffraction on the DMD $\vec{f_k}$ must equal $\vec{d}$, this leads to:

$$\begin{pmatrix} m & 0 & \frac{1}{2}(1-\cos\gamma+\cos\gamma) & \frac{1}{2}(1-\cos\gamma) & -ax \\ 0 & m & \frac{1}{2}(1-\cos\gamma) & \frac{1}{2}(1-\cos\gamma+\cos\gamma) & -ay \\ 0 & 0 & -\frac{1}{\sqrt{2}}\sin\gamma & \frac{1}{\sqrt{2}}\sin\gamma & -az \end{pmatrix} \begin{pmatrix} m_x \\ m_y \\ s \\ t \\ u \end{pmatrix} = \begin{pmatrix} h_x \\ h_y \\ h_z \end{pmatrix}$$

This equation system is under determined. To solve this, we need to check every mirror $(m_x, m_y)$, which leads to:

$$\begin{pmatrix} \frac{1}{2}(1-\cos\gamma+\cos\gamma) & \frac{1}{2}(1-\cos\gamma) & -ax \\ \frac{1}{2}(1-\cos\gamma) & \frac{1}{2}(1-\cos\gamma+\cos\gamma) & -ay \\ -\frac{1}{\sqrt{2}}\sin\gamma & \frac{1}{\sqrt{2}}\sin\gamma & -az \end{pmatrix} \begin{pmatrix} s \\ t \\ u \end{pmatrix} = \begin{pmatrix} h_x \\ h_y \\ h_z \end{pmatrix} - m \begin{pmatrix} m_x \\ m_y \\ 0 \end{pmatrix}$$

This has an analytic solution for each single mirror (see ray-dmd-intersection.nb in the attached repository). But there is only one valid Solution, which can be determined by checking the range of $s$ and $t$. If there are still more than one solution then the solution with the smallest $u$ is the valid one, because that is the position on the DMD which gets hit first.

**Analytic Phase Shifting Approach:**

The field of a single mirror can be written down as follows. The integral can be solved analytically. (see dmd.nb in the attached repository)

$$E^{single\ mirror}\left(\vec{a}, \vec{b}, \gamma^{\pm}_{m_x,m_y}, w, \lambda\right) = \iint_0^w \exp\left(\frac{2\pi i}{\lambda}\left(R\left(\gamma^{\pm}_{m_x,m_y}\right) \cdot (s,t,0)\right) \cdot \left(\vec{a} - \vec{b}\right)\right) ds\, dt$$

The field generated by the phase shift between the single mirrors can be written down as follows.

$$E^{shift}_{m_x,m_y}\left(\vec{a}, \vec{b}, m, \lambda\right) = \exp\left(\frac{2\pi i}{\lambda} \vec{m}(m_x, m_y) \cdot \left(\vec{a} - \vec{b}\right)\right)$$

**Grating Approach:**

The sum over the phase shifts of the single mirrors corresponds to the field of the native DMD grating. The sum can furthermore be simplified as follows.

$$E^{grating}(\vec{a},\vec{b},m,N_x,N_y,\lambda) = \sum_{m_x=0,m_y=0}^{N_x-1,N_y-1} E^{shift}_{m_x,m_y}(\vec{a},\vec{b},m,\lambda)$$
$$= \sum_{m_x=0,m_y=0}^{N_x-1,N_y-1} \exp\left(\frac{2\pi m i}{\lambda}\left((a_x-b_x)m_x+(a_y-b_y)m_y\right)\right)$$
$$= \frac{\left(-1+\exp\left(\frac{2\pi m i}{\lambda}(a_x-b_x)N_x\right)\right)\left(-1+\exp\left(\frac{2\pi m i}{\lambda}(a_y-b_y)N_y\right)\right)}{\left(-1+\exp\left(\frac{2\pi m i}{\lambda}(a_x-b_x)\right)\right)\left(-1+\exp\left(\frac{2\pi m i}{\lambda}(a_y-b_y)\right)\right)}$$

#### 2. Quick guide though the Coherent DMD Simulator Fiji-Plugin

After copying the file "coherent_dmd_sim_simulator.jar" into the ".../Fiji.app/plugins" directory, you can access our algorithms in Fiji under "Plugins>Coherent DMD Simulator>..." (Figure 1). The default settings of the plugin are chosen to reproduce the results in our article. Currently the plugin has no CUDA GPU support. If you want to use the GPU support, you must customize the .java files and start them from the command line or a development environment of your choice.

Figure 2 to Figure 5 show the input masks for the approaches presented in our article. Depending on the approach, you can specify various system constraints. The ray tracing approach Figure 2 and the analytic phase shifting approach Figure 3 each simulate one single diffraction pattern. The ability to simulate different patterns is currently not yet implemented in the ray tracing approach but will be available soon.

The grating approach simulates many diffraction images depending on the incidence parameters. For these the distance between the brightest diffraction order and the center of the envelope in pixels is determined. Additionally, the x and y position of the brightest diffraction order is determined. These data are stored e.g. at a simulation wavelength of 488 *nm* in the files "488_false-eveope-peak-distance.tif", "488_false-intensity-max-x.tif" and "488_false-intensity-max-y.tif". Before starting the grating approach, you will be asked whether the diffraction images should also be stored, which requires a lot of RAM. The results of the grating approach can be further processed using the diffraction space analysis (Figure 6) to find out under which diffraction angles a blaze condition is fulfilled.

All saved .tif files contain meta information about the system constraints set for the simulation. These can be displayed via "Image>Show Info" (Figure 7)

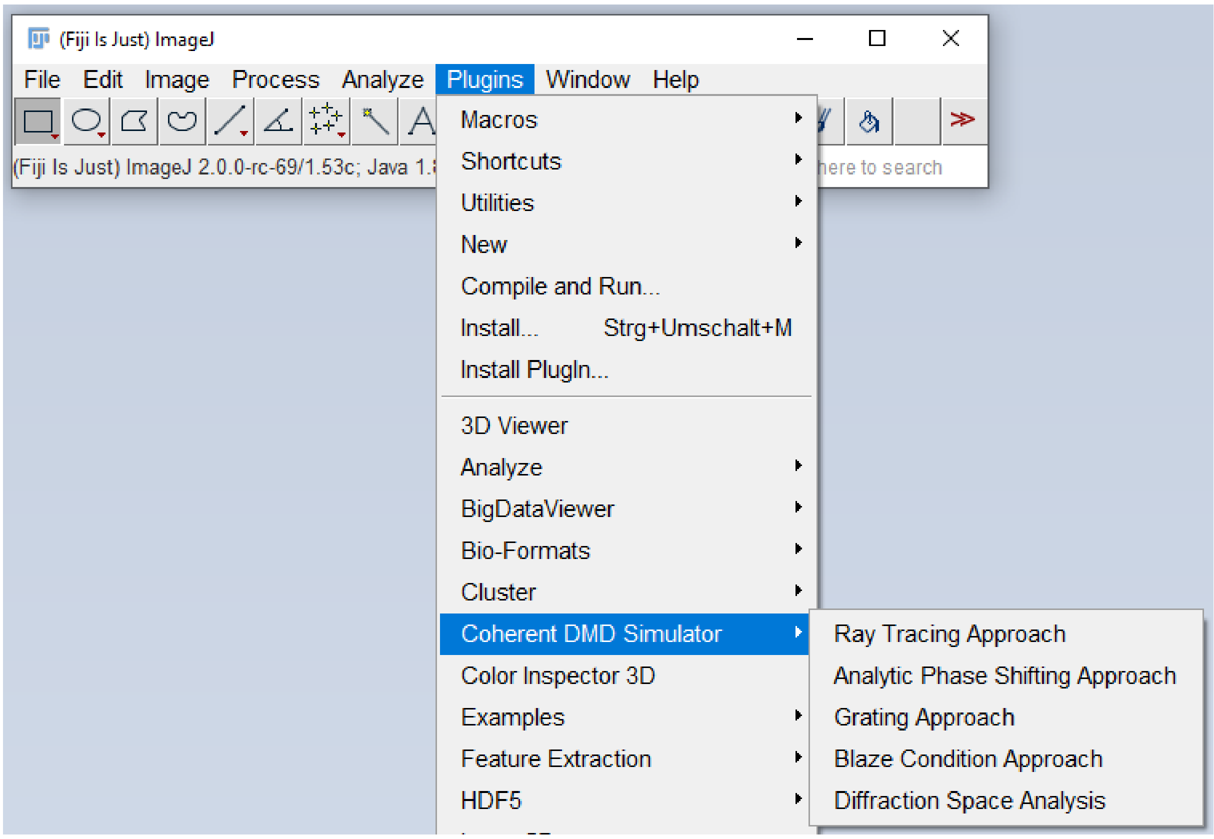


*Figure 1*

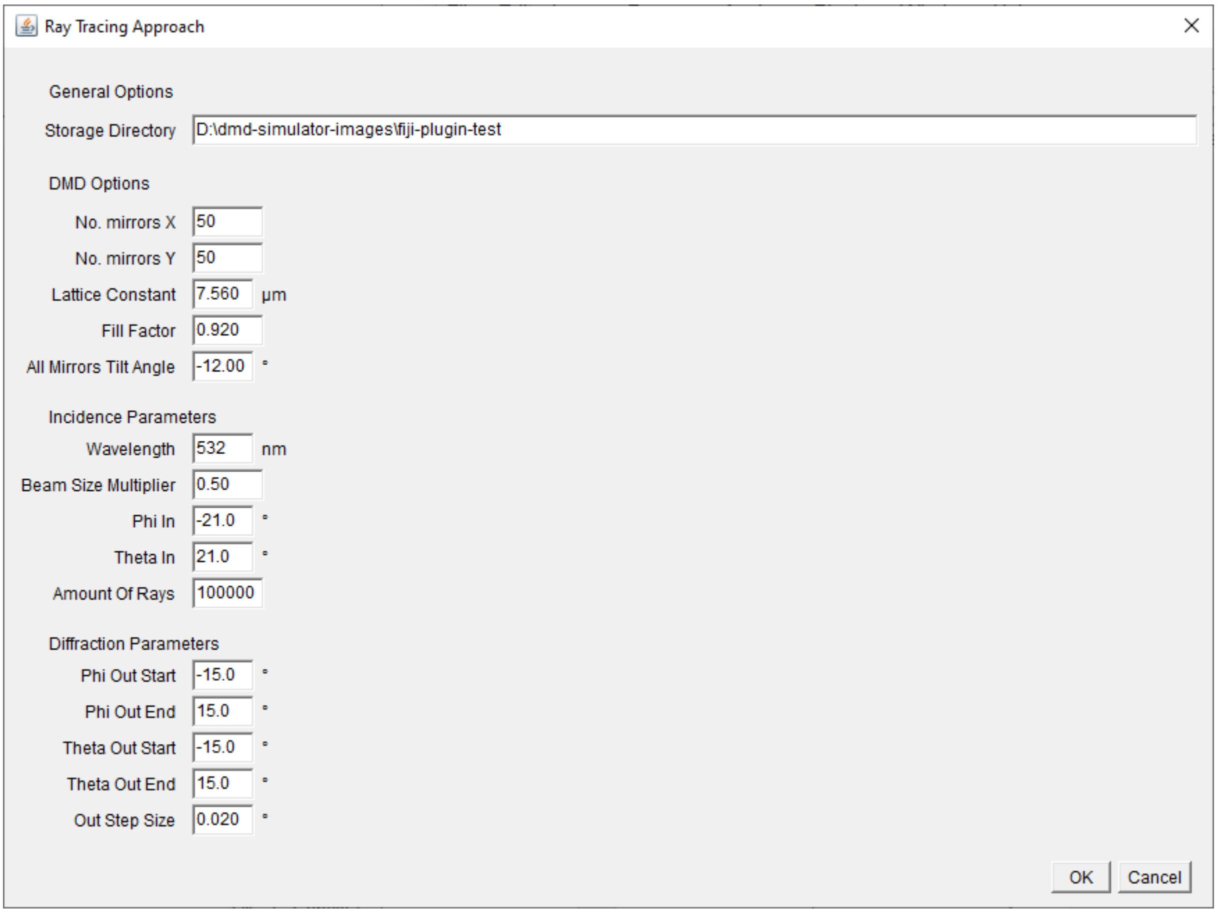


*Figure 2*

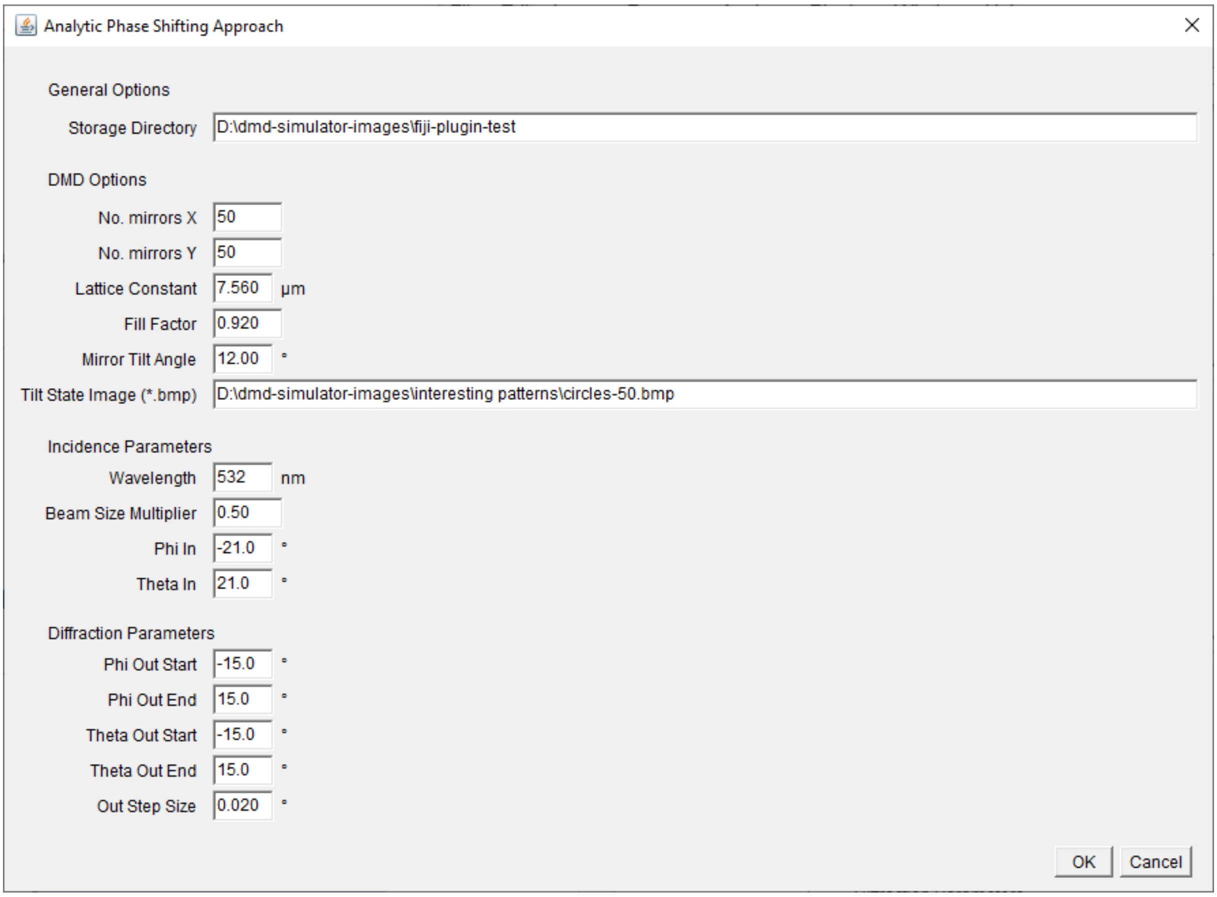
Analytic Phase Shifting Approach
General Options
Storage Directory
D:\dmd-simulator-images\fiji-plugin-test
DMD Options
No. mirrors X
50
No. mirrors Y
50
Lattice Constant
7.560
µm
Fill Factor
0.920
Mirror Tilt Angle
12.00
°
Tilt State Image (*.bmp)
D:\dmd-simulator-images\interesting patterns\circles-50.bmp
Incidence Parameters
Wavelength
532
nm
Beam Size Multiplier
0.50
Phi In
-21.0
°
Theta In
21.0
°
Diffraction Parameters
Phi Out Start
-15.0
°
Phi Out End
15.0
°
Theta Out Start
-15.0
°
Theta Out End
15.0
°
Out Step Size
0.020
°
OK
Cancel

*Figure 3*

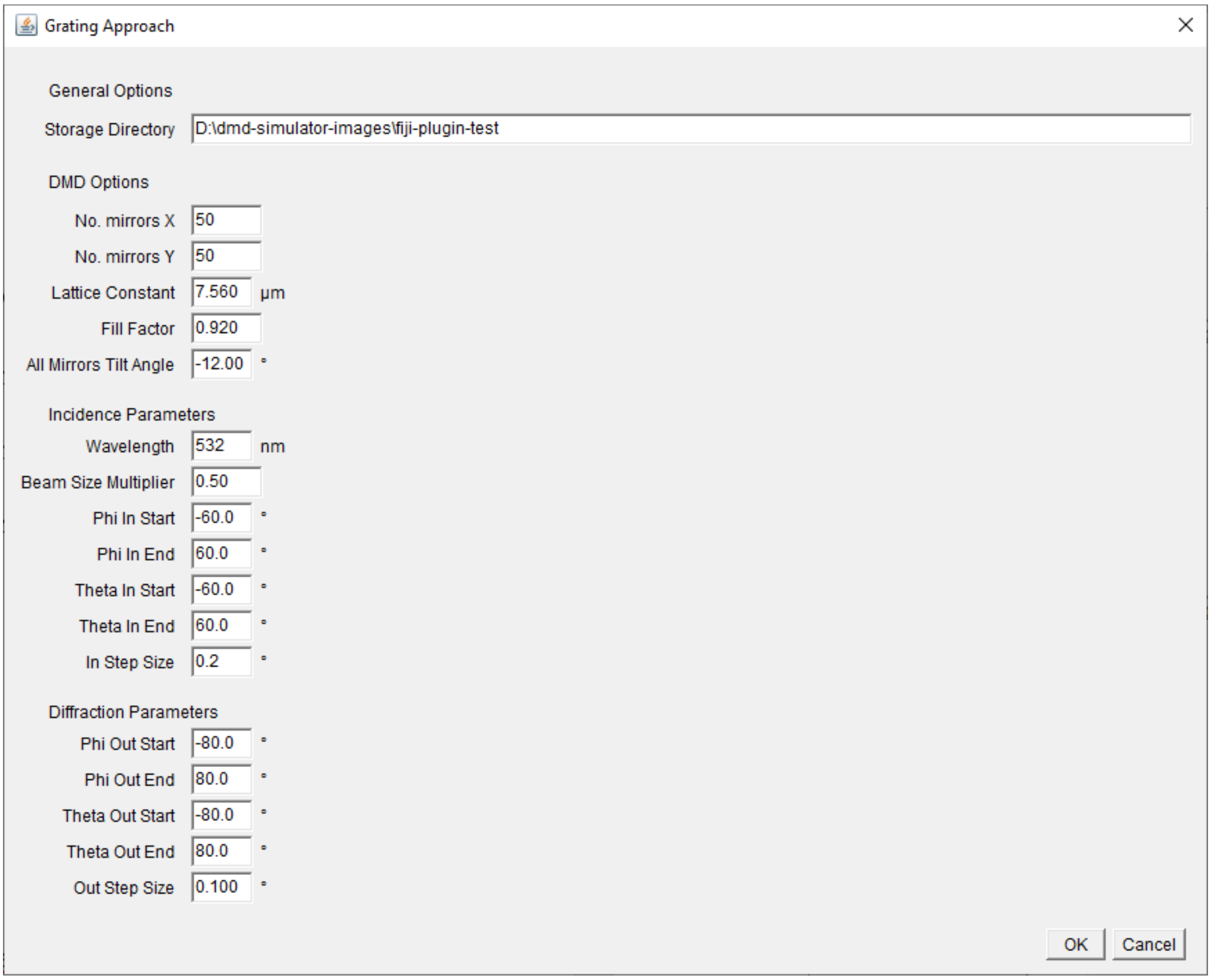


*Figure 4*

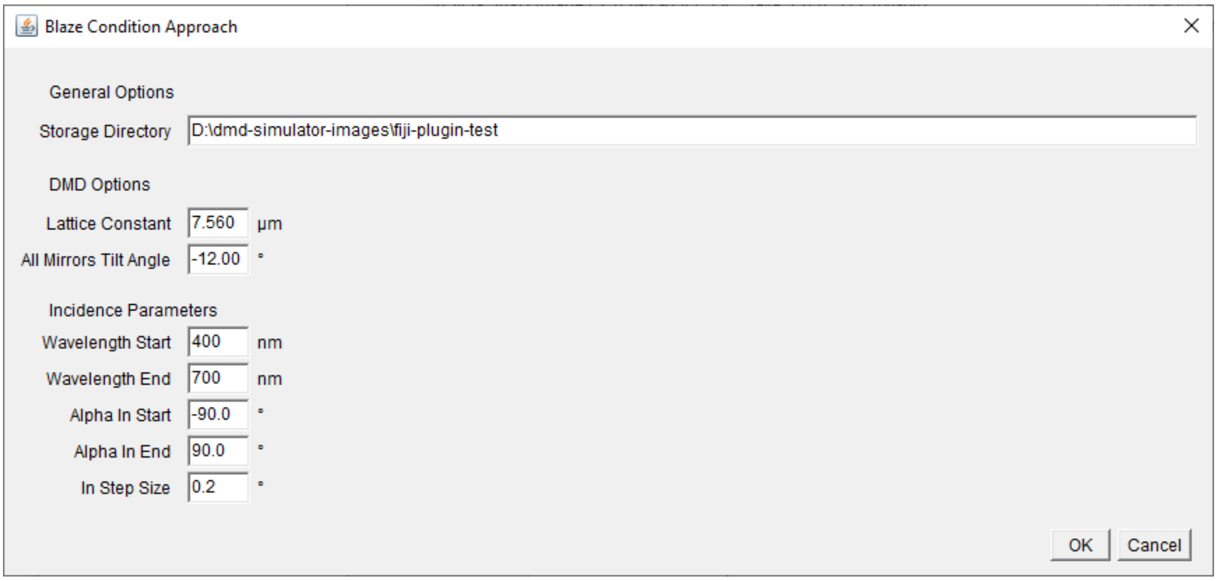


*Figure 5*

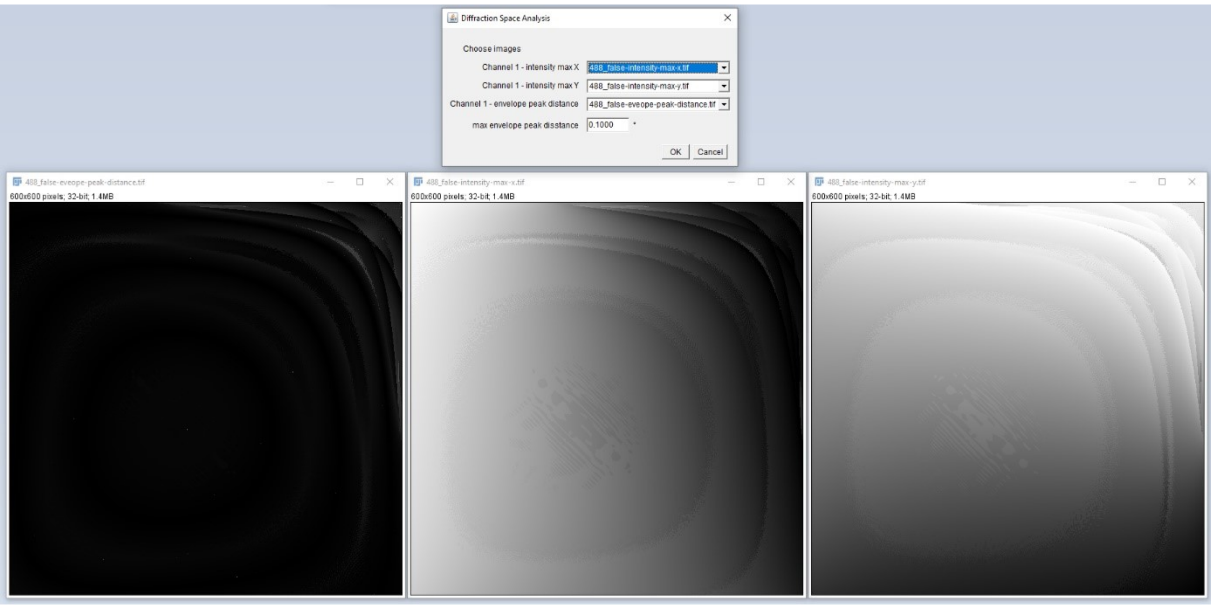


*Figure 6*

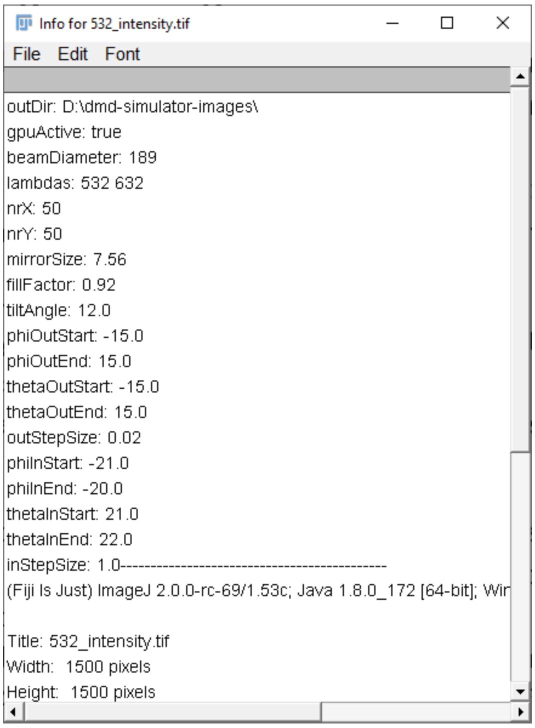


*Figure 7*
